# Huge narwhal displacement following shipping of iron ore in the Canadian Arctic Archipelago

**DOI:** 10.1101/2023.05.29.542701

**Authors:** Lars Witting

## Abstract

With 100,000 narwhals inhabiting populations in the Canadian Arctic Archipelago, the Northwest Passage runs through the largest aggregation of narwhals in the world. I use a metapopulation analysis that covers eight of the narwhal populations in the area to estimate that about 25,000 narwhals have emigrated from Eclipse Sound to Admiralty Inlet since 2007, causing almost total population collapse in Eclipse Sound following shipping of iron ore through Milne Inlet and Eclipse Sound. It is estimated that up to about 30,000 narwhals may vanish in the long run, unless shipping is increasingly regulated, or narwhals adapt to the disturbance and reinhabits the abandoned areas. Reflecting only a local disturbance, these population effects warn against intense shipping in other fjords with narwhals, and against a potentially much larger impact from increased shipping through the Northwest Passage in the future.

---

The Canadian Arctic Archipelago has the largest concentration of narwhals in the world, with estimates exceeding 100,000 individuals (Doniol-Valcroze et al. 2015). Until recently the area was largely undisturbed, but shipping activities have increased over the last three decades with a steep regime shift in overall shipping in 2007 (Pizzolato et al. 2014). This relates not only to traffic into the Northwest Passage at Lancaster Sound but even more so to intense shipping of iron ore from the Mary River Mine through Milne Inlet and Eclipse Sound (Kochanowicz et al. 2021).

Marine shipping disturb narwhals (Golder 2021; Tervo et al. 2021), but the long-term consequences for individuals and populations are unknown. Arctic odontocetes react to low ship noise (Finley et al. 1990; Halliday et al. 2020), with narwhals seen to react more than 40km from a controlled experiment even when sound exposure is below background noise (Tervo et al. 2021). Narwhal buzzing can halve at 12km, foraging can stop at 8km (Tervo et al. 2021), and individuals avoid ships at 4km (Golder 2020, 2021). Industrial impact assessments for narwhal assume that these responses are local in space and time, and that they do not accumulate to critical population effects (Golder 2021, 2022). Several population effects, however, are impossible to estimate from local disturbance studies. Rather than reacting to disturbance directly, a disturbance-imposed population displacements may follow as a gradual yearly change in the average migration of individuals, with critical population regulation effects emerging on even longer timescales.

I analyse for such accumulated population impacts on narwhals from increased shipping in Northeast Canada and Northwest Greenland. Individual narwhals have strong site fidelity migrating to the same summer foraging grounds year after year (Heide-Jørgensen et al. 2013), and I look for larger displacements among the eight summer feeding populations in the area (Fig. 1;

**Figure 1:**
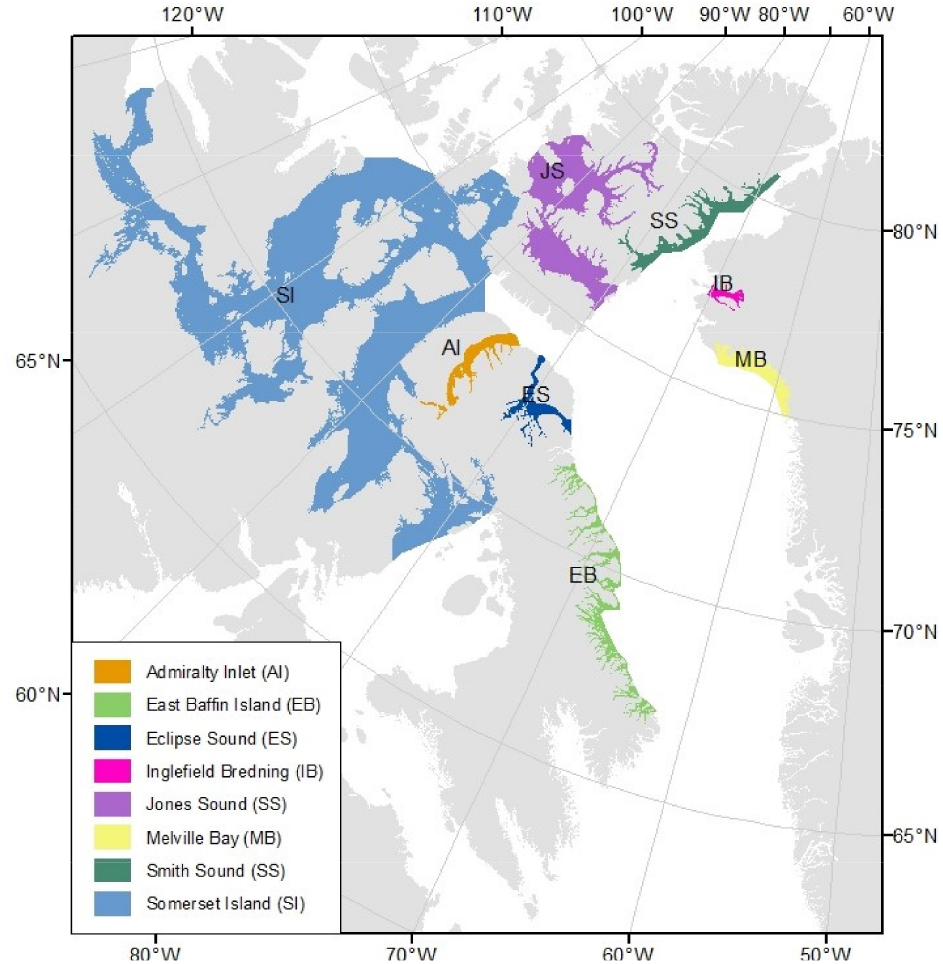
Summer distribution of narwhal populations in Northeast Canada and Northwest Greenland. From Watt et al. (2019)

Hobbs et al. 2019). I use the metapopulation model (Witting et al. 2019) developed by the JWG (2021), where catch allocation (Watt et al. 2019) links eight independent population dynamic models together. These are age-structured Bayesian models with density and selection regulation estimating the population dynamic trajectories from 1970 based on catch data and one to eight abundance estimates, dependent on population. I add two extra migration parameters to each population, allowing for independent emigration/immigration before and after the steep increase in shipping activity in 2007 (Pizzolato et al. 2014).

Using uniform migration priors that allow for up to 30% yearly emigration, and 30% yearly immigration, I obtain 16 posterior migration estimates across the eight populations (Fig. 2). All posterior migration estimates except two are either weakly updated by the Bayesian model or updated and centred around zero, showing absence of evidence for emigration and immigration. The two exceptions are Eclipse Sound after 2007 with a posterior annual emigration estimate of 13% (90% CI:9.4%–16%), and Admiralty Inlet after 2007 with a posterior annual immigration estimate of 5.9% (90% CI:3.0%–11%). With Admiralty Inlet and Eclipse Sound being neighbouring populations, this suggests substantial displacement of narwhals from Eclipse Sound to Admiralty Inlet.

**Figure 2:**
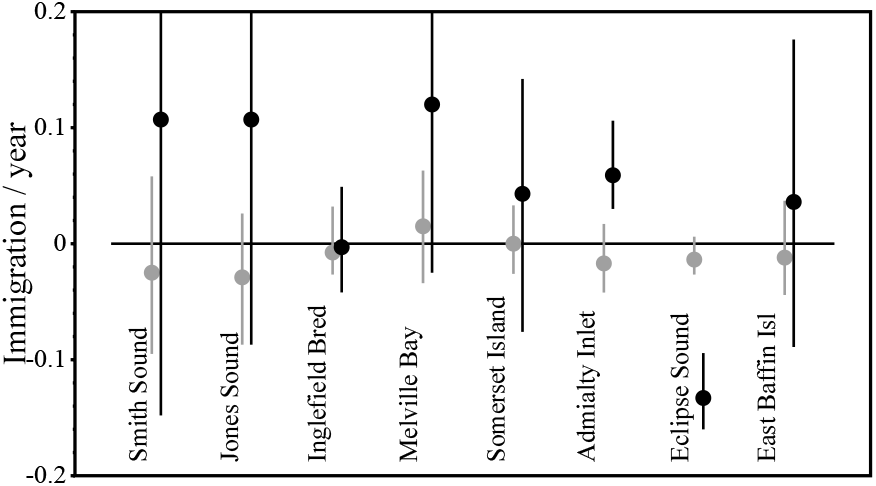
Posterior migration estimates for the eight narwhal summer populations, with independent estimates for periods before (gray) and after (black) 2007. Estimates are fractions of current abundance, with dots being medians and lines the 90% credibility intervals.

In the final model I exclude migration except for Eclipse Sound and Admiralty Inlet after 2007. Assuming a constant yearly migration of individuals, Fig. 3 shows the posterior population projections for Eclipse Sound and Admiralty Inlet. These are based on a posterior estimate of 1,540 (90% CI:1,040–2,240) narwhals emigrating annually from Eclipse Sound from 2007 to 2022, and 1,720 (90% CI:927–2,380) narwhals immigrating annually to Admiralty Inlet. A total of 24,640 (90% CI:16,640–35,840) narwhals have so far emigrated from the Eclipse Sound population, with a corresponding total of 27,520 (90% CI:14,830–38,080) narwhals immigrating into Admiralty Inlet from 2007 to 2022. With a posterior population estimate of 203 (90% CI:0– 1,170) narwhals in 2022, it is estimated that there will be almost no narwhals left in Eclipse Sound in 2023.

**Figure 3:**
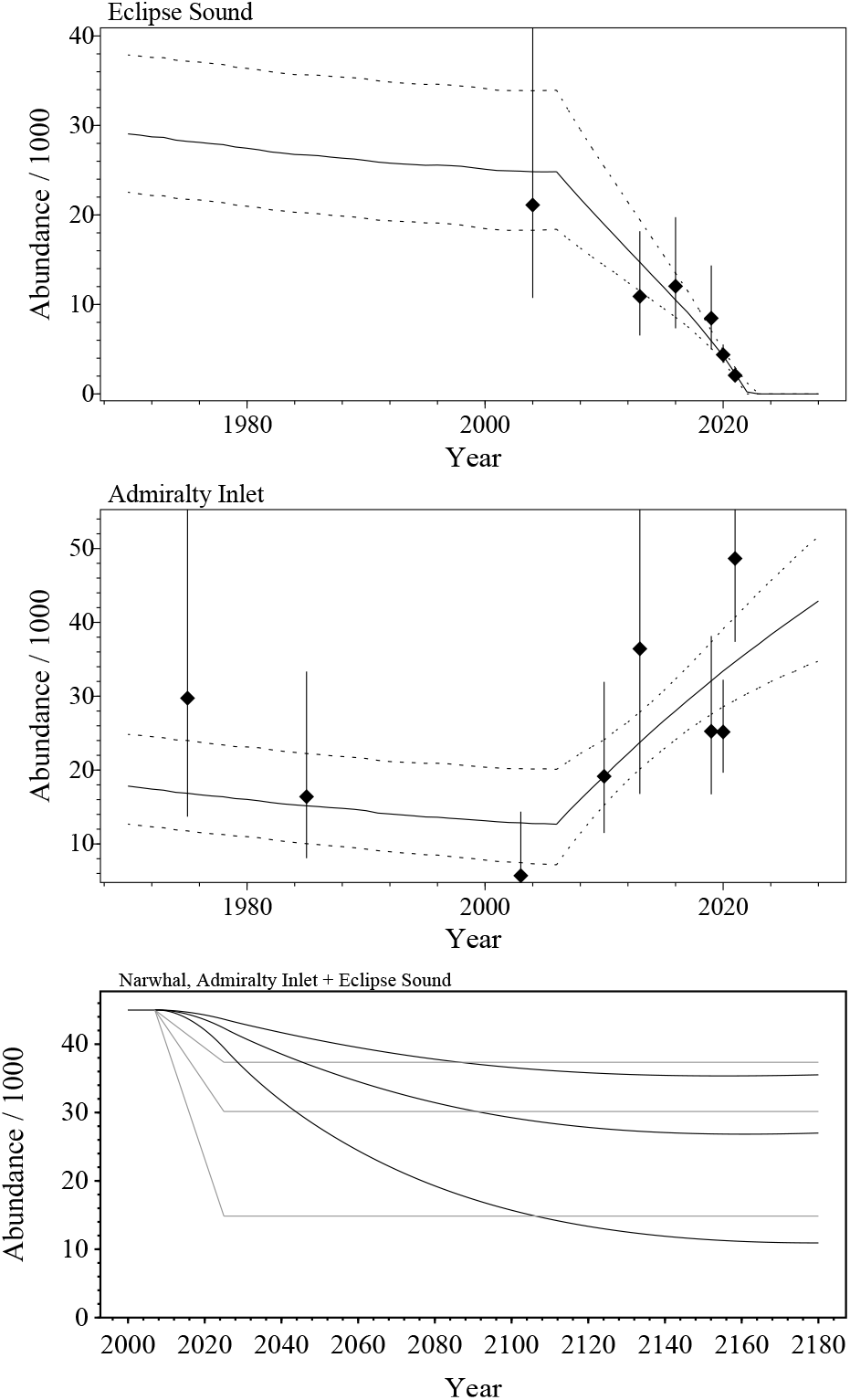
Narwhal population trajectories for Eclipse Sound and Admiralty Inlet. Points with bars are abundance estimates with 90% CI, solid curves the median of the estimated trajectories, and dotted curves the 90% credibility intervals. Bottom plot: Assuming a joint abundance of 45,000 narwhals for Admiralty Inlet and Eclipse Sound by 2025 and a historical abundance of 15,000, black curves are the expected trajectories should 1*/*6, 1*/*2 or the complete population regulation occur during August and September, with grey curves being the corresponding equilibria from the estimated shift in distribution.

Having pronounced preferences for sea ice and cold Arctic water, narwhals are considered sensitive to climatic changes (Laidre et al. 2008; Heide-Jørgensen et al. 2020). Yet, the observed displacement is not in line with climate changes in sea surface temperature (SST). While mean SST during the open water season (August and September) in Eclipse Sound was lower in the 1990s (−0.1°C) than during the migration period from 2007 to 2022 (1.5°C), a climate response is not expected as SST declined in Eclipse Sound and Admiralty Inlet during the migration period (Chambault et al. 2020; Fig. 4; Table S5). During the first half of the period (from 2007 to 2014) where the displacement started, SST was even lower in Eclipse Sound (2.1°C) than in Admiralty Inlet (2.8°C), and it was higher along East Baffin Island (3.2°C) where no emigration was observed between surveys in 2003 and 2013.

**Figure 4:**
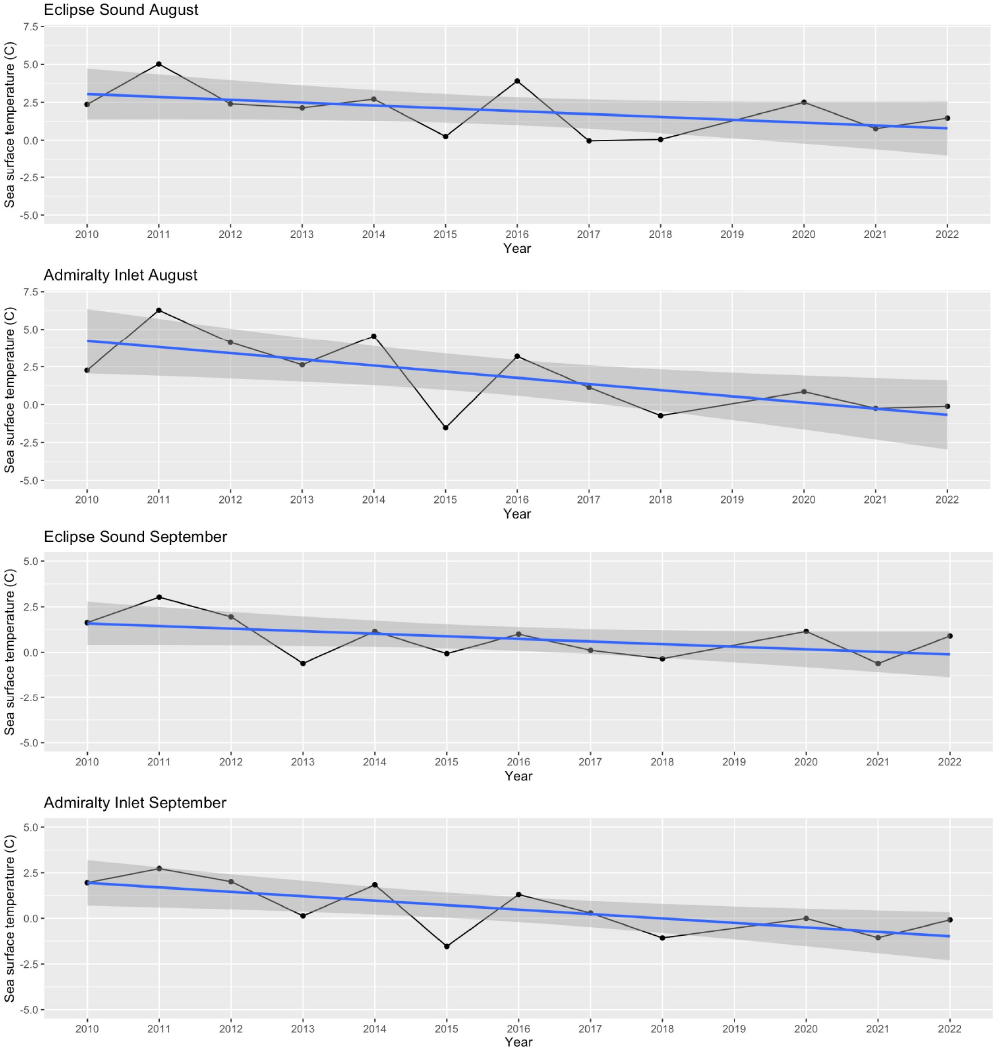
Displacement by climate change is not supported by the observed decline since 2010 in the mean surface temperature in Eclipse Sound and Admiralty Inlet in August and September. Dots are data obtained from the Global Ocean Physics Reanalysis Glorys S2V4 (PHYS-001-024) and the Global Ocean Physics Reanalysis Glorys12v1 (PHY-001-030), see Chambault et al. (2020) for details. Lines and grey areas are linear regressions with 95% CI.

Other potential hypotheses for the displacement include distributional shifts in predators and prey (Golder 2022). Killer whales e.g. have been established in the Canadian Arctic for centuries with a sporadic and likely increasing summer occurrence, including Admiralty Inlet and Eclipse Sound (Higdon et al. 2012; Lefort et al. 2020). But, during a simultaneous tracking of killer whales and narwhals in Admiralty Inlet, the tracked narwhals did not leave the inlet but moved instead to shallow coastal waters when killer whales were present (Breed et al. 2017). Quite generally, there seems also to be an absence of evidence for a shift in the distribution of predators and prey between Admiralty Inlet and Eclipse Sound.

The collapse of the Eclipse Sound population, however, follows the increase in ship traffic in the area. Not only is no migration estimated by the metapopulation model prior to the steep traffic increase in 2007 (Pizzolato et al. 2014), but the migratory response after 2007 is in agreement with area specific differences in ship traffic. Kochanowicz et al. (2021) used Automatic Identification System data from 2015 to 2018 to identify areas with ship noise above 120dB, which may negatively impact marine mammal hearing and behaviour (NMFS 2016; Gomez et al. 2016). Eclipse Sound and Milne Inlet have by far the highest frequency of exposures above 120dB, following the increased shipment of iron ore from the Mary River Mine (Kochanowicz et al. 2021). With two ore carrier vessel transits per day during the open water season in 2020 (Golder 2021), no other fjord system with narwhals in Northeast Canada and Northwest Greenland have similar amounts of heavy ship traffic. The second highest, and much lower, frequency of exposures above 120dB is found in Lancaster Sound and Parry Chanel, and no exposures above 120dB were estimated for the majority of Admiralty Inlet (Kochanowicz et al. 2021).

I agree with JDW (2022) that the increased shipping of iron ore is by far the most likely cause for the huge—an almost complete—displacement of narwhals from Eclipse Sound. Unless shipping is increasingly regulated, or narwhals adapt to the disturbance and reinhabit the abandoned areas, we may expect longterm consequences that extend far beyond the currently observed lumping of narwhals. The metapopulation model estimates the equilibrium abundance of narwhals in Admiralty Inlet to 17,800 (90% CI:12,700–24,800) individuals. While narwhal populations are likely partially regulated by winter feeding, if summer regulation is essential, we cannot exclude that up to about 30,000 narwhals may vanish (Fig. 3, bottom plot). Reflecting only a local disturbance, these population effects warn not only against substantial shipping in other fjords with narwhals, but also against a potentially much larger metapopulation impact caused by increased shipping through the Northwest Passage in the future.

## Acknowledgements

I thank Philippine Chambault for compiling sea surface temperatures, and Mads Peter Heide-Jørgensen for comments.

## A Supplementary material/methods

I use the metapopulation model (Witting et al. 2019) developed by the JWG (2021), where catch allocation (Watt et al. 2019) links eight independent population dynamic models together. These are age-structured Bayesian models that estimate *i*) the population dynamic trajectories from 1970 based on catch data and the one to eight abundance estimates available per population (Table S1), and *ii*) the median (*m*) and first (*m*_0_) age of reproductive maturity from maturity data. The population models are regulated by density and selection following Witting (2021), and I add two extra migration parameters to each model to allow for independent emigration/immigration before and after the steep increase in shipping activity in 2007 (Pizzolato et al. 2014).

The log likelihood ln *L* = ln *L*_*n*_ + ln *L*_*m*_ of a population model is calculated by the joint likelihood of the abundance data (*L*_*n*_) for that population and data on the fraction of mature females at given ages (*L*_*m*_, data from Garde 2020). The method of de la Mare (1986) is used to calculate the likelihood component of the abundance data under the assumption that observation errors are log-normally distributed (Buckland 1992)

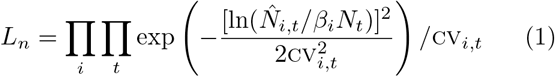

where 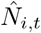 is the point estimate of the *i*th set of abundance data in year *t*, cv_*i,t*_ is the coefficient of variation of the estimate, *N*_*t*_ is the simulated abundance, and *β*_*i*_ a bias term with is set to one for absolute abundance estimates.

The likelihood component on the age of reproductive maturity is assumed multinomially distributed

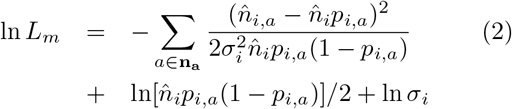

where 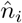 is the total number of females with an estimated age of maturity, 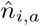 is the number of these females that mature at age *a, p*_*i,a*_ is the simulated probability that a female would mature at age *a*, and *σ*_*i*_ is potential overdispersal (no overdispersion assumed here, i.e., *σ*_*i*_ = 1). More details on the mathematics, statistics, and biological reasoning of the model are described by Witting et al. (2019).

This supplementary information lists the abundance estimates in Table S1, the prior distributions of the model parameters in Table S2, the sampling resampling statistics of the Bayesian integration in Table S4, and the posterior parameter estimates in Table S3. Estimates of sea surface temperature are listed in Table S5. The realised prior and posterior distributions of the fitted parameters are shown for the eight population models in Fig. S5 to Fig. S12. Model fits to the age of maturity data are shown in Fig. S13, and the estimated trajectories in Fig. S14.

**Table 1:** Abundance estimates with CVs for summer aggregations of narwhal. Data from JWG (2021), Golder (2021, 2022), and JDW (2022)

| Year | Smith | Jones | Inglefield | Melville | Somerset | Admiralty | Eclipse | Baffin |
| --- | --- | --- | --- | --- | --- | --- | --- | --- |
| 1975 | - | - | - | - | - | 29740; 0.47 | - | - |
| 1981 | - | - | - | - | 32520; 0.22 | - | - | - |
| 1985 | - | - | 3690; 0.32 | - | - | 16400; 0.43 | - | - |
| 1986 | - | - | 9560; 0.4 | - | - | - | - | - |
| 1996 | - | - | - | - | 45360; 0.35 | - | - | - |
| 2001 | - | - | 3010; 0.41 | - | - | - | - | - |
| 2002 | - | - | 1940; 0.33 | - | 29770; 0.58 | - | - | - |
| 2003 | - | - | - | - | - | 5700; 0.56 | - | 10710; 0.34 |
| 2004 | - | - | - | - | - | - | 21110; 0.41 | - |
| 2007 | - | - | 4110; 0.29 | 1830; 0.94 | - | - | - | - |
| 2009 | - | - | - | - | - | - | - | - |
| 2010 | - | - | - | - | - | 19160; 0.31 | - | - |
| 2012 | - | - | - | 920; 0.48 | - | - | - | - |
| 2013 | 17010; 0.68 | 13200; 0.38 | - | - | 51730; 0.28 | 36430; 0.47 | 10900; 0.31 | 11990; 0.4 |
| 2014 | - | - | - | 1770; 0.5 | - | - | - | - |
| 2016 | - | - | - | - | - | - | 12040; 0.3 | - |
| 2019 | - | - | 2870; 0.28 | 4760; 0.77 | - | 25260; 0.25 | 8460; 0.32 | - |
| 2020 | - | - | - | - | - | 25170; 0.15 | 4380; 0.14 | - |
| 2021 | - | - | - | - | - | 48650; 0.16 | 2080; 0.17 | - |

**Table 2:** Priors for the parameters of the different models (*M*). *N* the abundance, *c* the removals, ϑ the female fraction at birth, *γ* the density regulation, *γ*_*ι*_ the inertia, *i*_0_ the initial life history, *a*_*m*_ the rep. maturity, *m*_0_ the first rep. maturity, *n* the migration, *p*_0_ the first year survival, *p* the yearly survival, and *\** denotes population dynamic equilibrium. Abundance is given in thousands. The prior probability distribution is given by superscripts; *p*: fixed value, *u*: uniform (min;max), *U* : log uniform (min;max), *b*: beta 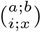 with *i*=min and *x*=max, and *f* : distribution from file.

| $M$ | $N_0$ | $N^{**}$ | $c$ | $\vartheta$ | $\gamma$ | $\gamma_i$ | $i_0$ | $a_m$ | $m_0$ | $n$ | $p_0$ | $p$ |
| --- | --- | --- | --- | --- | --- | --- | --- | --- | --- | --- | --- | --- |
| smi | - | 2;80<sup>U</sup> | f | .5<sup>p</sup> | .95<sup>p</sup> | .69<sup>p</sup> | 1<sup>p</sup> | 9.7;10<sup>u</sup> | 9.2;10<sup>u</sup> | - | .5;1<sup>u</sup> | 2;2<sup>b</sup><br>.95;1 |
| jon | - | 4;40<sup>U</sup> | f | .5<sup>p</sup> | .95<sup>p</sup> | .69<sup>p</sup> | 1<sup>p</sup> | 9.7;10<sup>u</sup> | 9.2;10<sup>u</sup> | - | .5;1<sup>u</sup> | 2;2<sup>b</sup><br>.95;1 |
| ing | 3;10<sup>U</sup> | 3;15<sup>U</sup> | f | .5<sup>p</sup> | .95<sup>p</sup> | .69<sup>p</sup> | 2;2<sup>b</sup><br>.5;1.5 | 9.7;10<sup>u</sup> | 9.2;10<sup>u</sup> | - | .5;1<sup>u</sup> | 2;2<sup>b</sup><br>.95;1 |
| mel | 1.2;10<sup>U</sup> | 2.2;15<sup>U</sup> | f | .5<sup>p</sup> | .95<sup>p</sup> | .69<sup>p</sup> | 2;2<sup>b</sup><br>.5;1.5 | 9.7;10<sup>u</sup> | 9.2;10<sup>u</sup> | - | .5;1<sup>u</sup> | 2;2<sup>b</sup><br>.95;1 |
| som | 10;50<sup>U</sup> | 30;90<sup>U</sup> | f | .5<sup>p</sup> | .95<sup>p</sup> | .69<sup>p</sup> | 2;2<sup>b</sup><br>.5;1.5 | 9.7;10<sup>u</sup> | 9.2;10<sup>u</sup> | - | .5;1<sup>u</sup> | 2;2<sup>b</sup><br>.95;1 |
| ecl | - | 15;50<sup>U</sup> | f | .5<sup>p</sup> | .95<sup>p</sup> | .69<sup>p</sup> | 1<sup>p</sup> | 9.7;10<sup>u</sup> | 9.2;10<sup>u</sup> | -3000;0<sup>u</sup> | .5;1<sup>u</sup> | 2;2<sup>b</sup><br>.95;1 |
| adm | - | 5;35<sup>U</sup> | f | .5<sup>p</sup> | .95<sup>p</sup> | .69<sup>p</sup> | 1<sup>p</sup> | 9.7;10<sup>u</sup> | 9.2;10<sup>u</sup> | 0;3000<sup>u</sup> | .5;1<sup>u</sup> | 2;2<sup>b</sup><br>.95;1 |
| baf | - | 5;30<sup>U</sup> | f | .5<sup>p</sup> | .95<sup>p</sup> | .69<sup>p</sup> | 1<sup>p</sup> | 9.7;10<sup>u</sup> | 9.2;10<sup>u</sup> | - | .5;1<sup>u</sup> | 2;2<sup>b</sup><br>.95;1 |

**Table 3:** Parameter estimates for the different models (*M*). Estimates are given by the median (*x*_.5_) and the 90% credibility interval (*x*_.05_ - *x*_.95_) of the posterior distributions. *N* the abundance, *b* the birth rate, *c* the removals, *d* the depletion ratio, *i*_0_ the initial life history, *a*_*m*_ the rep. maturity, *m*_0_ the first rep. maturity, *n* the migration, *p*_0_ the first year survival, *p* the yearly survival, and *\** denotes population dynamic equilibrium. Year by subscript: t:2022. Abundance is given in thousands.

| $M$ | | $N_0$ | $N_t$ | $N^{**}$ | $b^{**}$ | $c$ | $d_t$ | $i_0$ | $a_m$ | $m_0$ | $n$ | $p_0$ | $p$ |
| --- | --- | --- | --- | --- | --- | --- | --- | --- | --- | --- | --- | --- | --- |
| smi | $x_{.5}$ | - | 16.5 | 17.4 | .101 | .117 | .959 | 1 | 10.1 | 9.77 | - | .756 | .972 |
| | $x_{.05}$ | - | 5.61 | 6.31 | .031 | .020 | .846 | 1 | 9.91 | 9.48 | - | .525 | .957 |
| | $x_{.95}$ | - | 47.2 | 48.1 | .207 | .491 | .988 | 1 | 10.2 | 9.89 | - | .979 | .989 |
| jon | $x_{.5}$ | - | 12.7 | 13.6 | .096 | .112 | .94 | 1 | 10.1 | 9.77 | - | .762 | .973 |
| | $x_{.05}$ | - | 7.02 | 7.73 | .030 | .015 | .864 | 1 | 9.91 | 9.49 | - | .523 | .957 |
| | $x_{.95}$ | - | 23.1 | 23.9 | .206 | .446 | .972 | 1 | 10.2 | 9.88 | - | .976 | .989 |
| ing | $x_{.5}$ | 5.63 | 2.65 | 8.06 | .083 | .082 | .316 | .982 | 10.1 | 9.76 | - | .753 | .976 |
| | $x_{.05}$ | 3.74 | 1.68 | 5.78 | .028 | .017 | .168 | .598 | 9.92 | 9.48 | - | .527 | .959 |
| | $x_{.95}$ | 7.5 | 3.83 | 13.6 | .191 | .386 | .556 | 1.34 | 10.2 | 9.88 | - | .977 | .99 |
| mel | $x_{.5}$ | 3.86 | 1.24 | 7.56 | .109 | .074 | .159 | 1 | 10.1 | 9.76 | - | .748 | .971 |
| | $x_{.05}$ | 2.39 | .405 | 4.86 | .038 | .023 | .047 | .656 | 9.91 | 9.47 | - | .524 | .956 |
| | $x_{.95}$ | 6.21 | 2.68 | 13.7 | .221 | .307 | .376 | 1.37 | 10.2 | 9.89 | - | .98 | .987 |
| som | $x_{.5}$ | 28.8 | 50.5 | 59.2 | .109 | .582 | .867 | 1.04 | 10.1 | 9.76 | - | .759 | .97 |
| | $x_{.05}$ | 17.6 | 34.8 | 39.7 | .044 | .138 | .588 | .681 | 9.91 | 9.49 | - | .529 | .956 |
| | $x_{.95}$ | 43.6 | 77.7 | 84.8 | .202 | .817 | 1.19 | 1.38 | 10.2 | 9.89 | - | .972 | .985 |
| ecl | $x_{.5}$ | - | .203 | 29.1 | .106 | .403 | .007 | 1 | 10.1 | 9.78 | -1540 | .741 | .971 |
| | $x_{.05}$ | - | 0 | 22.5 | .034 | .143 | 0 | 1 | 9.9 | 9.45 | -2240 | .527 | .956 |
| | $x_{.95}$ | - | 1.17 | 37.9 | .212 | .636 | .047 | 1 | 10.2 | 9.89 | -1040 | .97 | .989 |
| adm | $x_{.5}$ | - | 35 | 17.8 | .093 | .387 | 1.98 | 1 | 10.1 | 9.77 | 1720 | .753 | .973 |
| | $x_{.05}$ | - | 29.3 | 12.7 | .030 | .117 | 1.27 | 1 | 9.92 | 9.48 | 927 | .524 | .956 |
| | $x_{.95}$ | - | 41.5 | 24.8 | .205 | .708 | 2.93 | 1 | 10.2 | 9.89 | 2380 | .974 | .989 |
| baf | $x_{.5}$ | - | 10.1 | 13.7 | .104 | .343 | .754 | 1 | 10.1 | 9.77 | - | .739 | .971 |
| | $x_{.05}$ | - | 6.48 | 9.81 | .034 | .081 | .622 | 1 | 9.92 | 9.47 | - | .523 | .955 |
| | $x_{.95}$ | - | 15.7 | 19.2 | .217 | .698 | .845 | 1 | 10.2 | 9.89 | - | .974 | .989 |

**Table 4:** Sampling statistics for the different models (*M*). The number of parameter sets in the sample (*n*_*S*_) and the resample (*n*_*R*_), the number of unique parameter sets in the resample, and the maximum number of occurrences of a unique parameter set in the resample. *n*_*S*_ and *n*_*R*_ are given in thousands.

| $M$ | $n_S$ | $n_R$ | unique | max |
| --- | --- | --- | --- | --- |
| smi | 300 | 1 | 968 | 3 |
| jon | 300 | 1 | 976 | 3 |
| ing | 300 | 1 | 858 | 5 |
| mel | 300 | 1 | 850 | 4 |
| som | 300 | 1 | 945 | 3 |
| ecl | 800 | 2 | 1086 | 14 |
| adm | 400 | 2 | 1669 | 6 |
| baf | 300 | 1 | 963 | 3 |

**Table 5:**
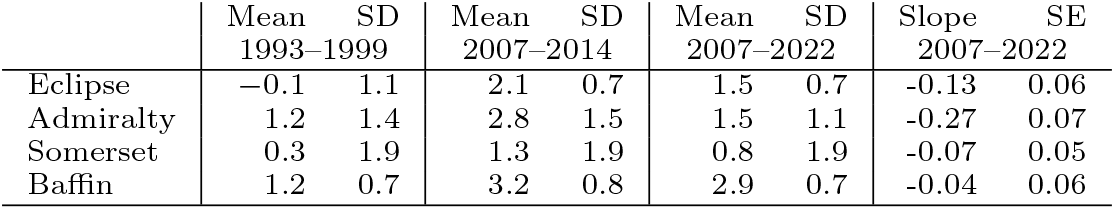
Sea surface temperature in the open water season (August and September) on four narwhal summering grounds, including the slope of the linear regression for the period 2007 to 2022. Data collected by Philippine Chambault from the Global Ocean Physics Reanalysis Glorys S2V4 (PHYS-001-024) and the Global Ocean Physics Reanalysis Glorys12v1 (PHY-001-030), see Chambault et al. (2020) for details.

|  | Mean<br>1993–1999 | SD<br>1993–1999 | Mean<br>2007–2014 | SD<br>2007–2014 | Mean<br>2007–2022 | SD<br>2007–2022 | Slope<br>2007–2022 | SE<br>2007–2022 |
| --- | --- | --- | --- | --- | --- | --- | --- | --- |
| Eclipse | -0.1 | 1.1 | 2.1 | 0.7 | 1.5 | 0.7 | -0.13 | 0.06 |
| Admiralty | 1.2 | 1.4 | 2.8 | 1.5 | 1.5 | 1.1 | -0.27 | 0.07 |
| Somerset | 0.3 | 1.9 | 1.3 | 1.9 | 0.8 | 1.9 | -0.07 | 0.05 |
| Baffin | 1.2 | 0.7 | 3.2 | 0.8 | 2.9 | 0.7 | -0.04 | 0.06 |

**Figure 5:**
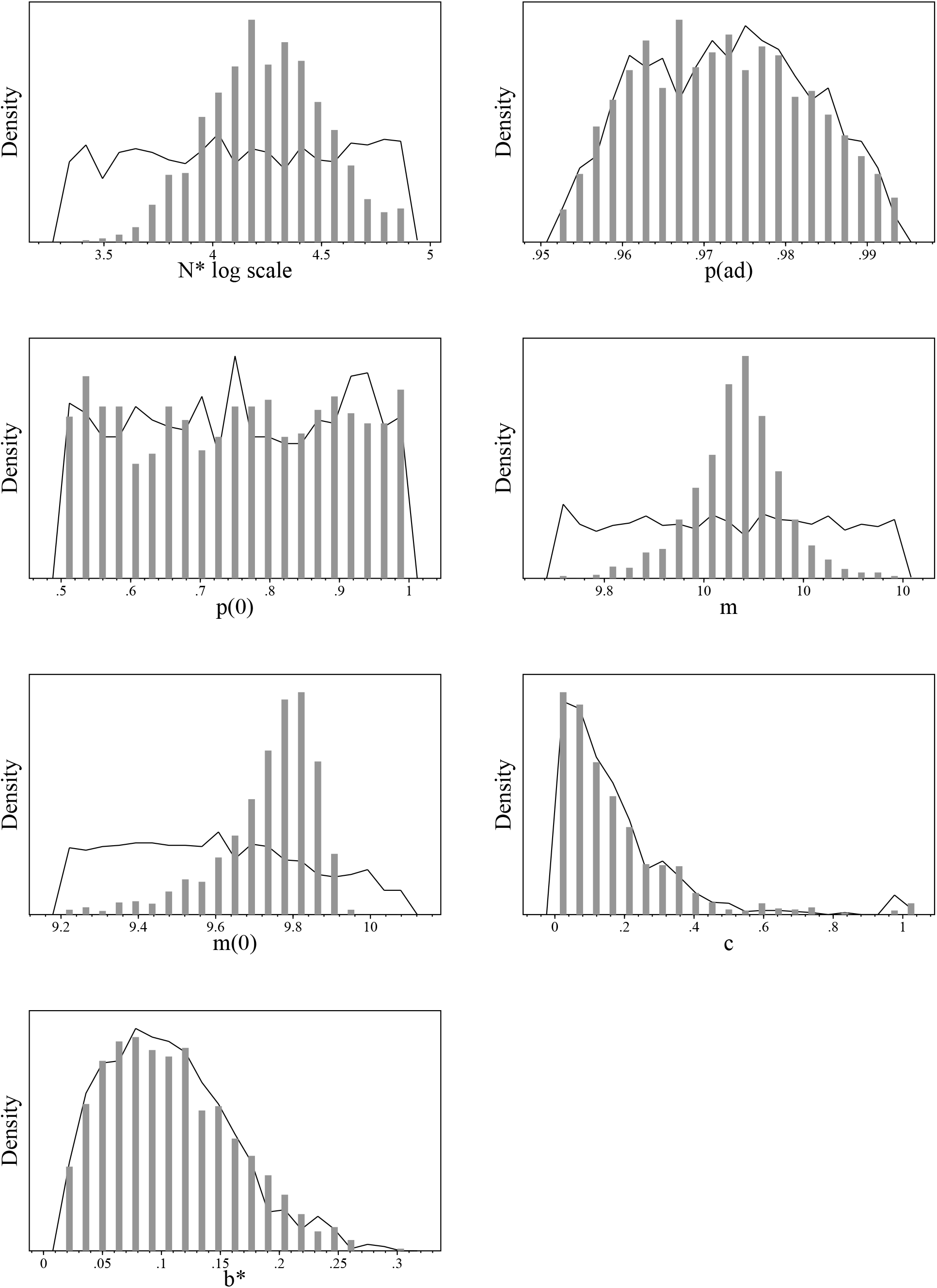
Smith Sound. Realised prior (curve) and posterior (bars) distributions.

**Figure 6:**
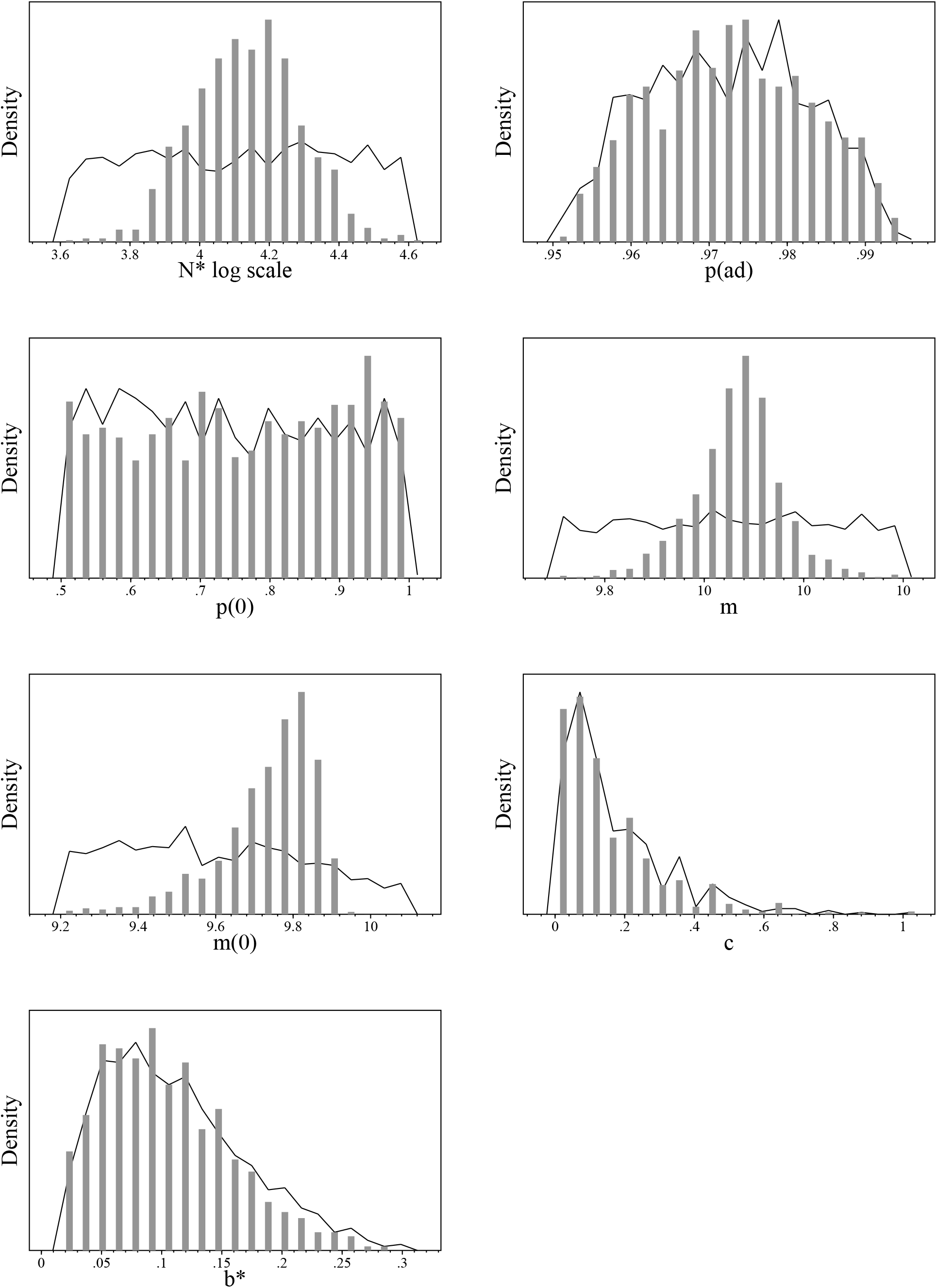
Jones Sound. Realised prior (curve) and posterior (bars) distributions.

**Figure 7:**
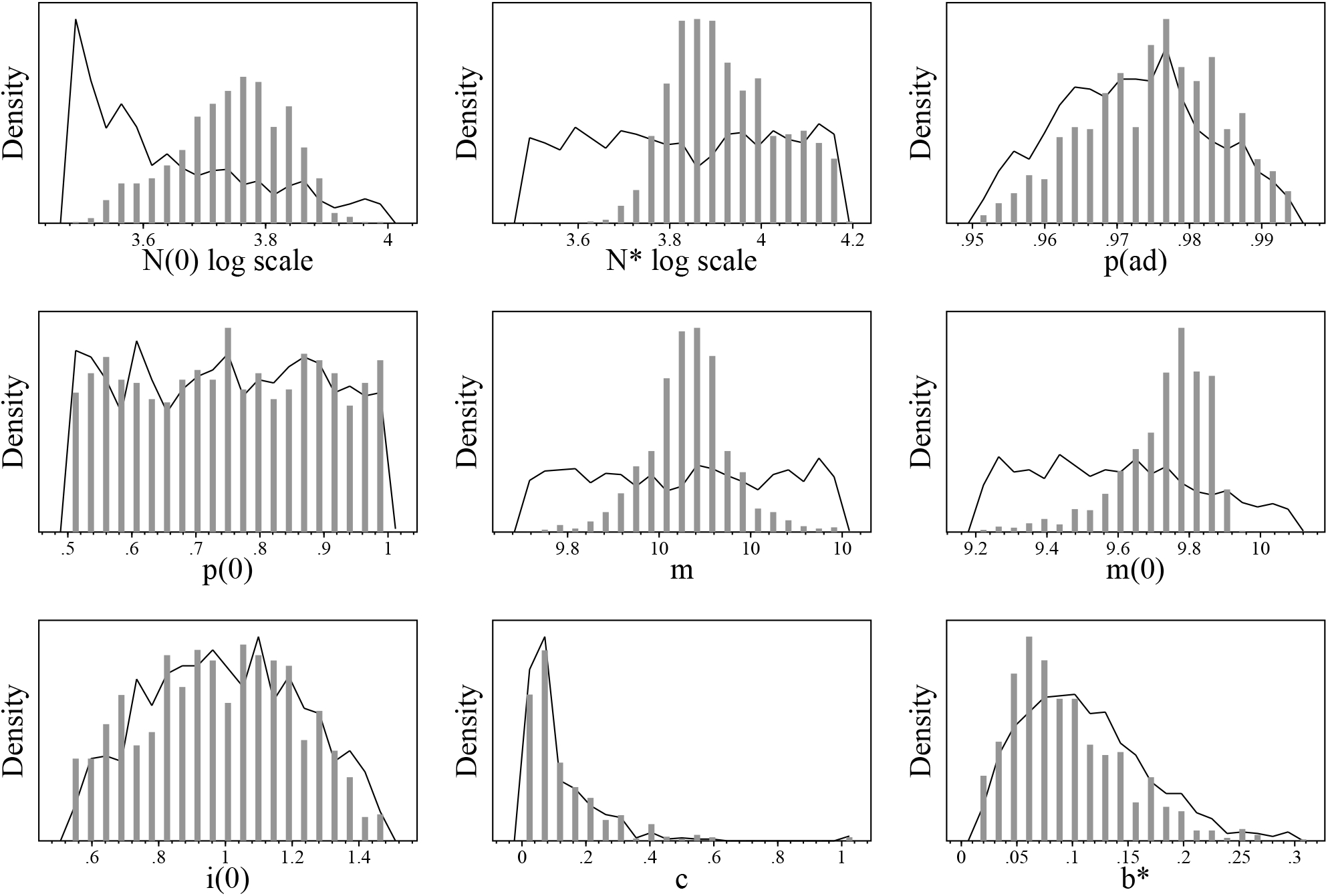
Inglefield Bredning. Realised prior (curve) and posterior (bars) distributions.

**Figure 8:**
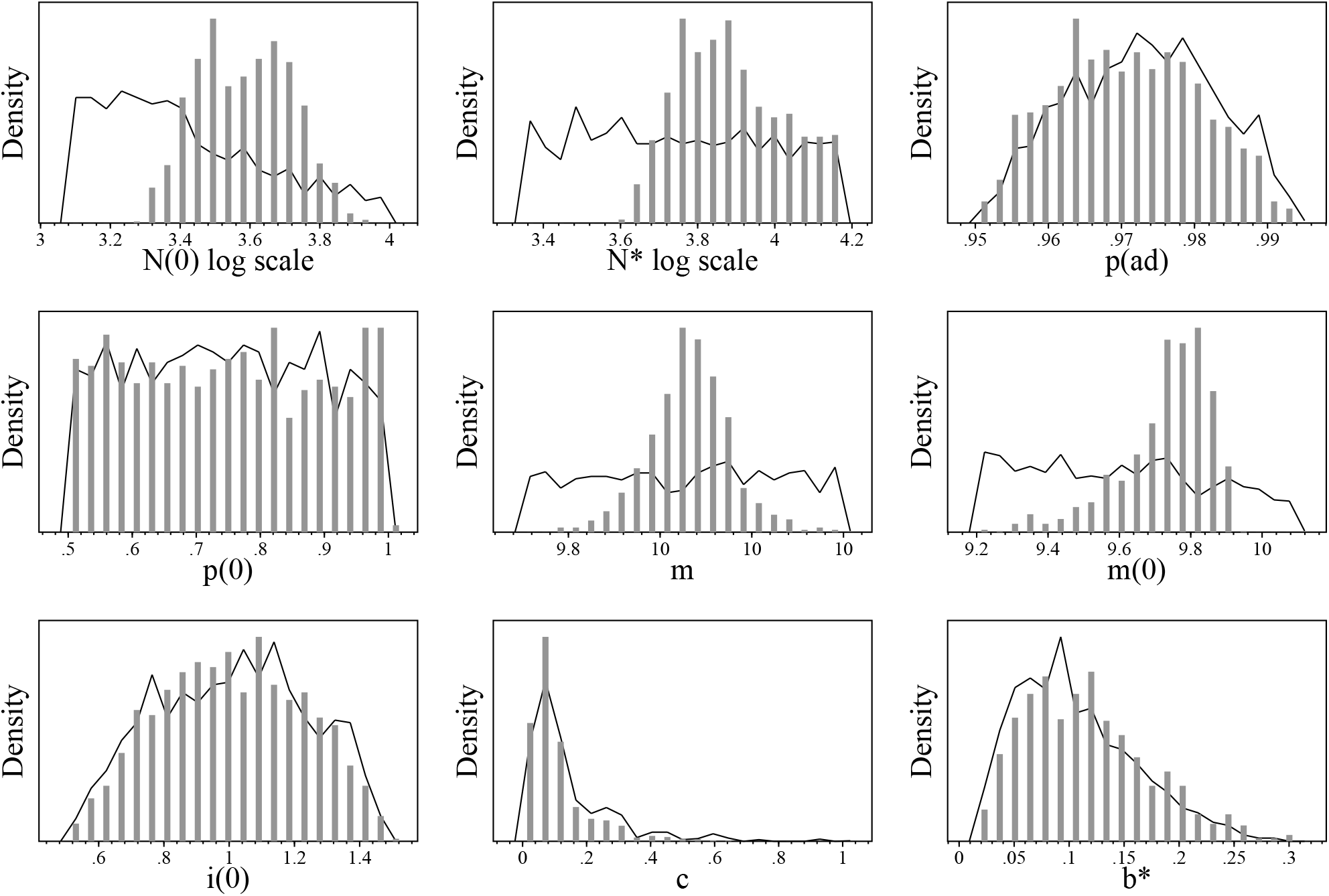
Melville Bay. Realised prior (curve) and posterior (bars) distributions.

**Figure 9:**
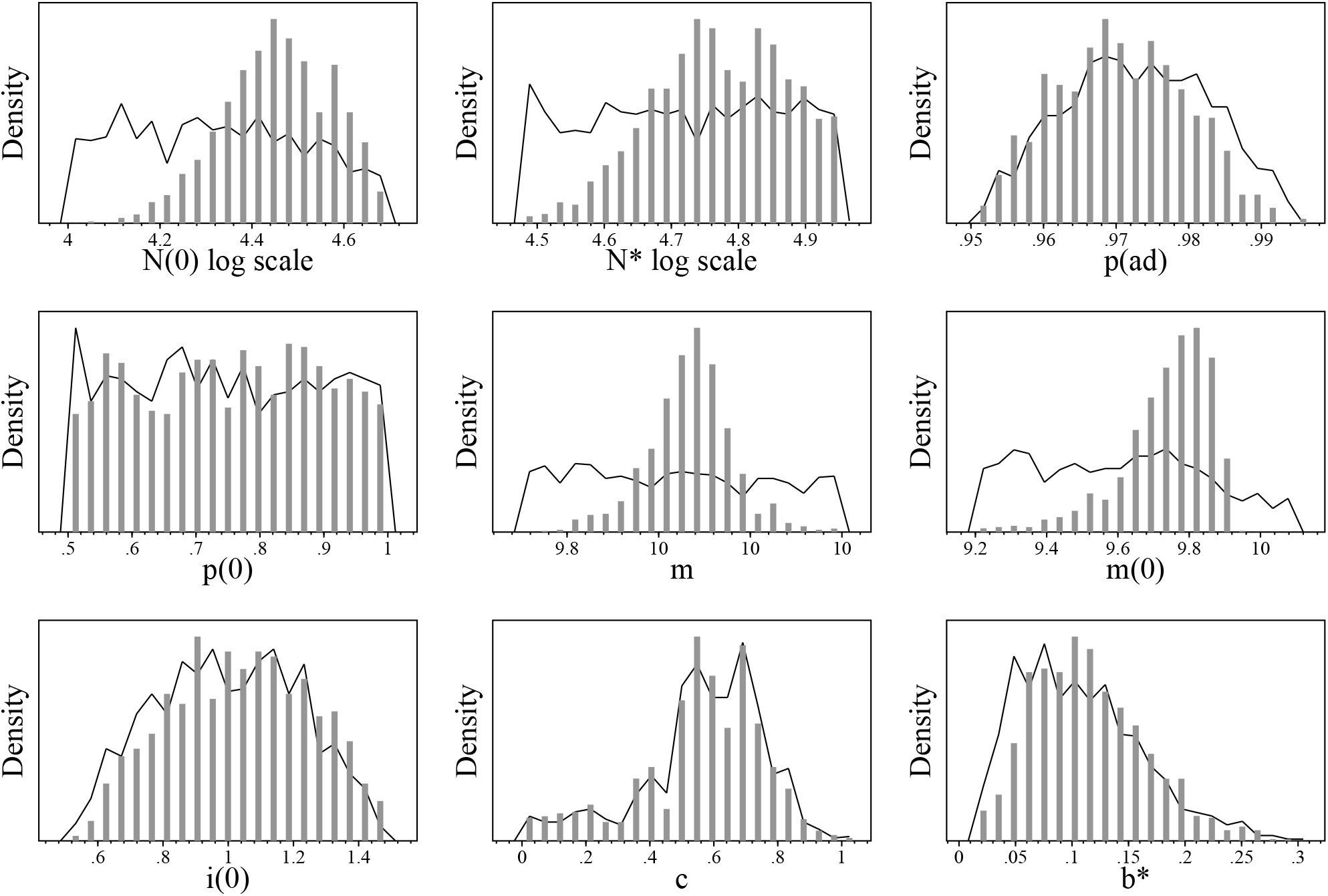
Somerset Island. Realised prior (curve) and posterior (bars) distributions.

**Figure 10:**
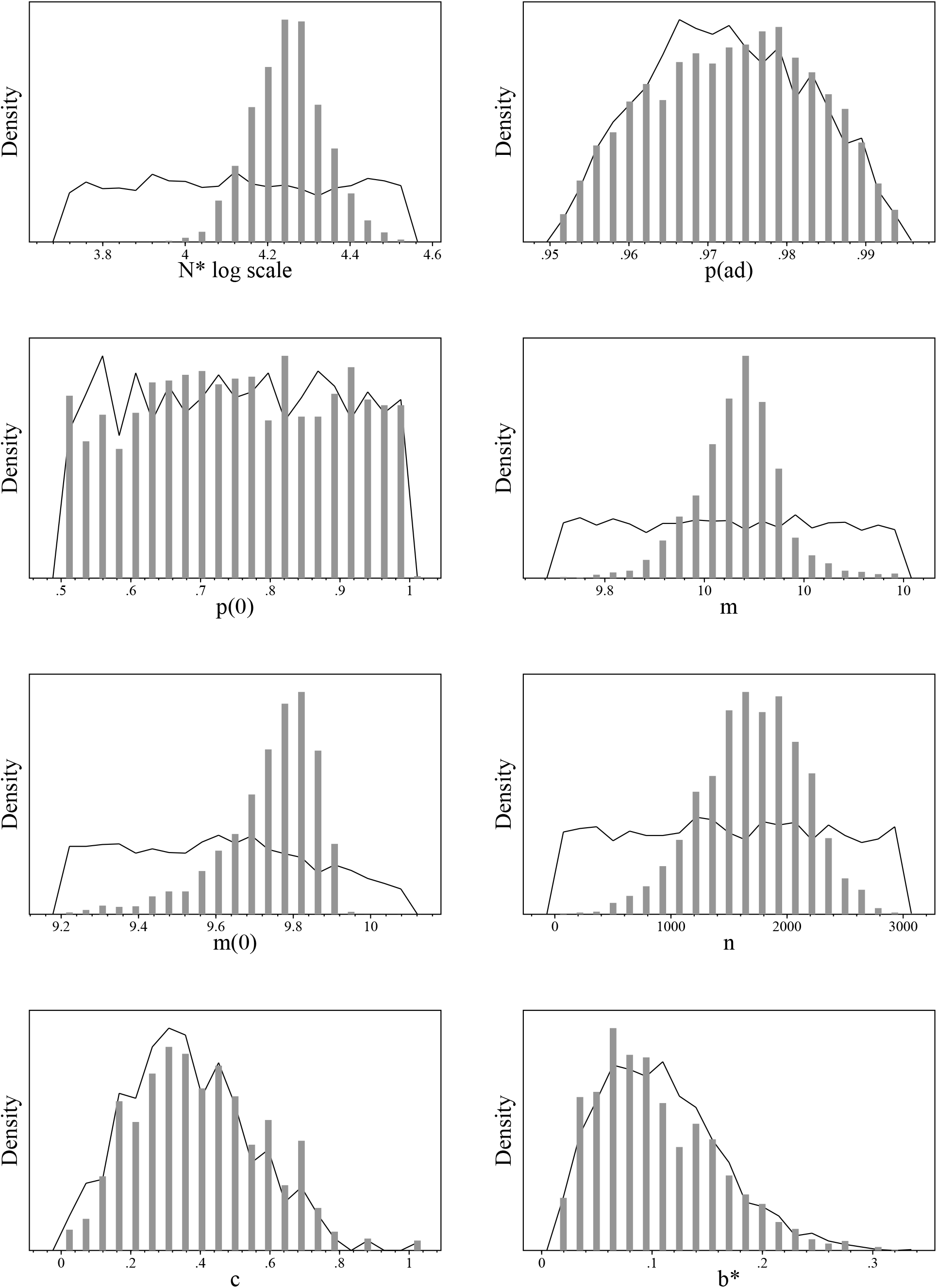
Admiralty Inlet. Realised prior (curve) and posterior (bars) distributions.

**Figure 11:**
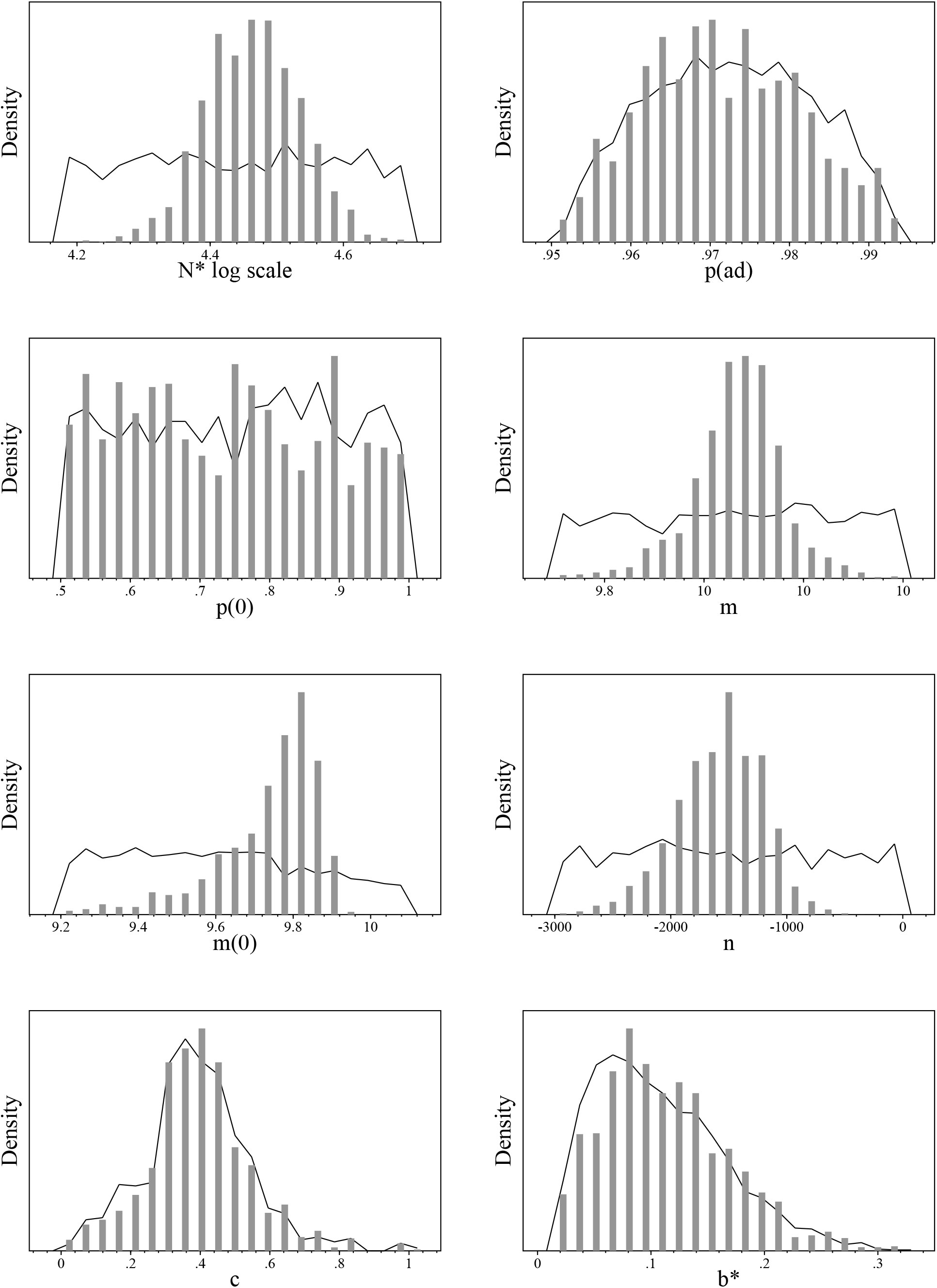
Eclipse Sound. Realised prior (curve) and posterior (bars) distributions.

**Figure 12:**
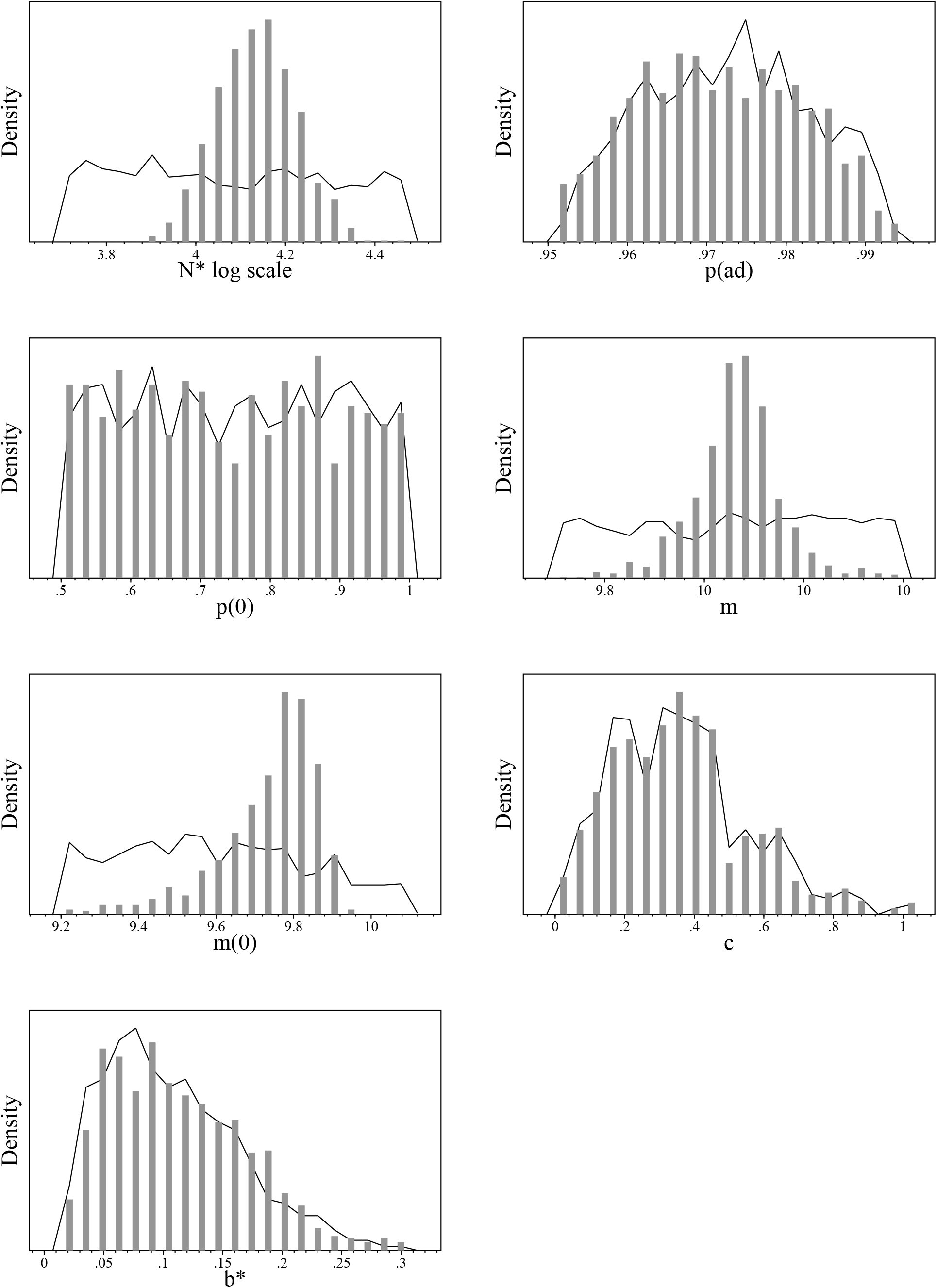
East Baffin Island. Realised prior (curve) and posterior (bars) distributions.

**Figure 13:**
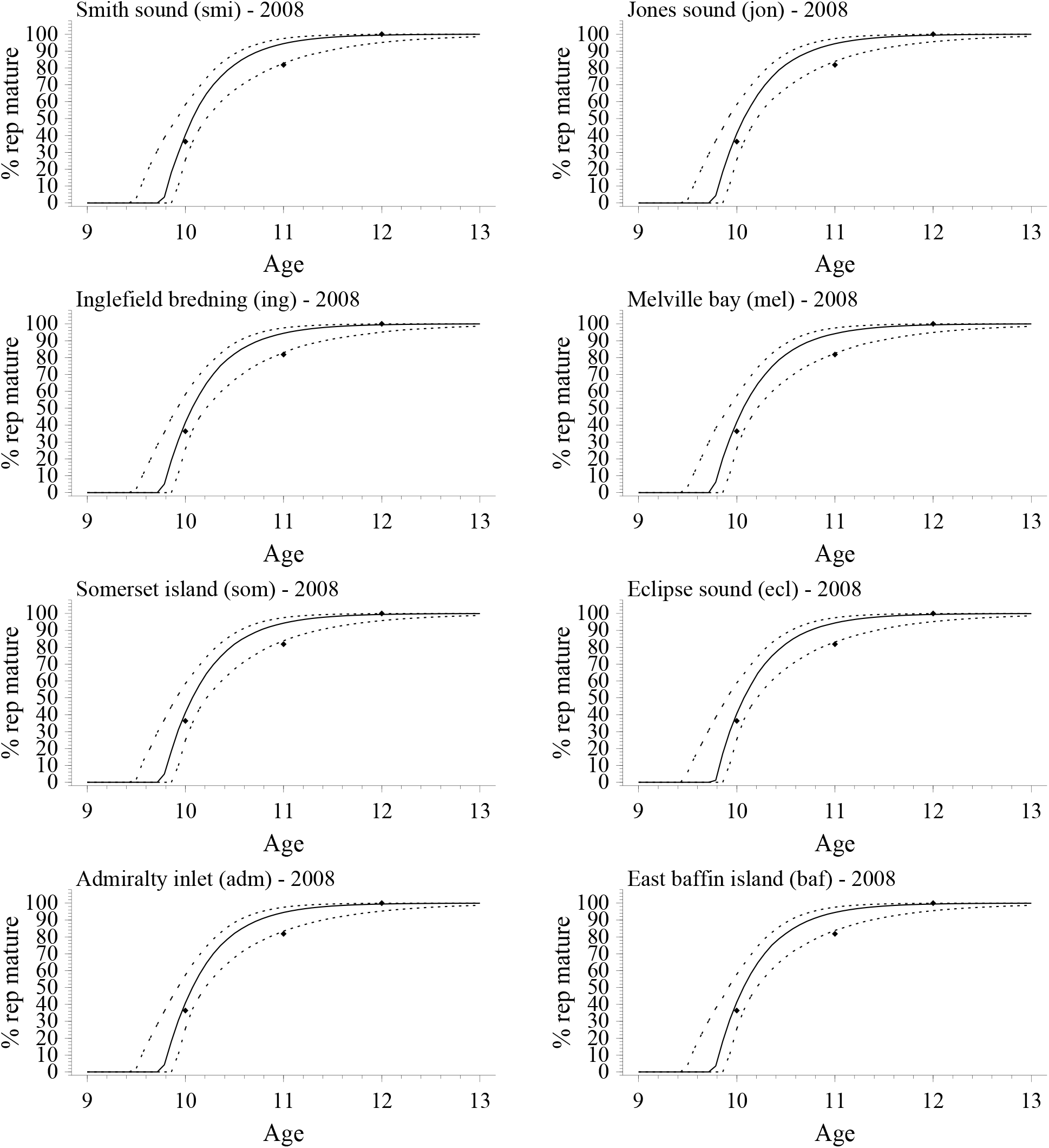
Reproductive maturity. The fraction of females that have reproduced at least once as a function of age. The solid and dashed curves are the median and 90% CI of the estimated models, and the points are the fraction among 11 female narwhals from Greenland (data from Garde, 2020).

**Figure 14:**
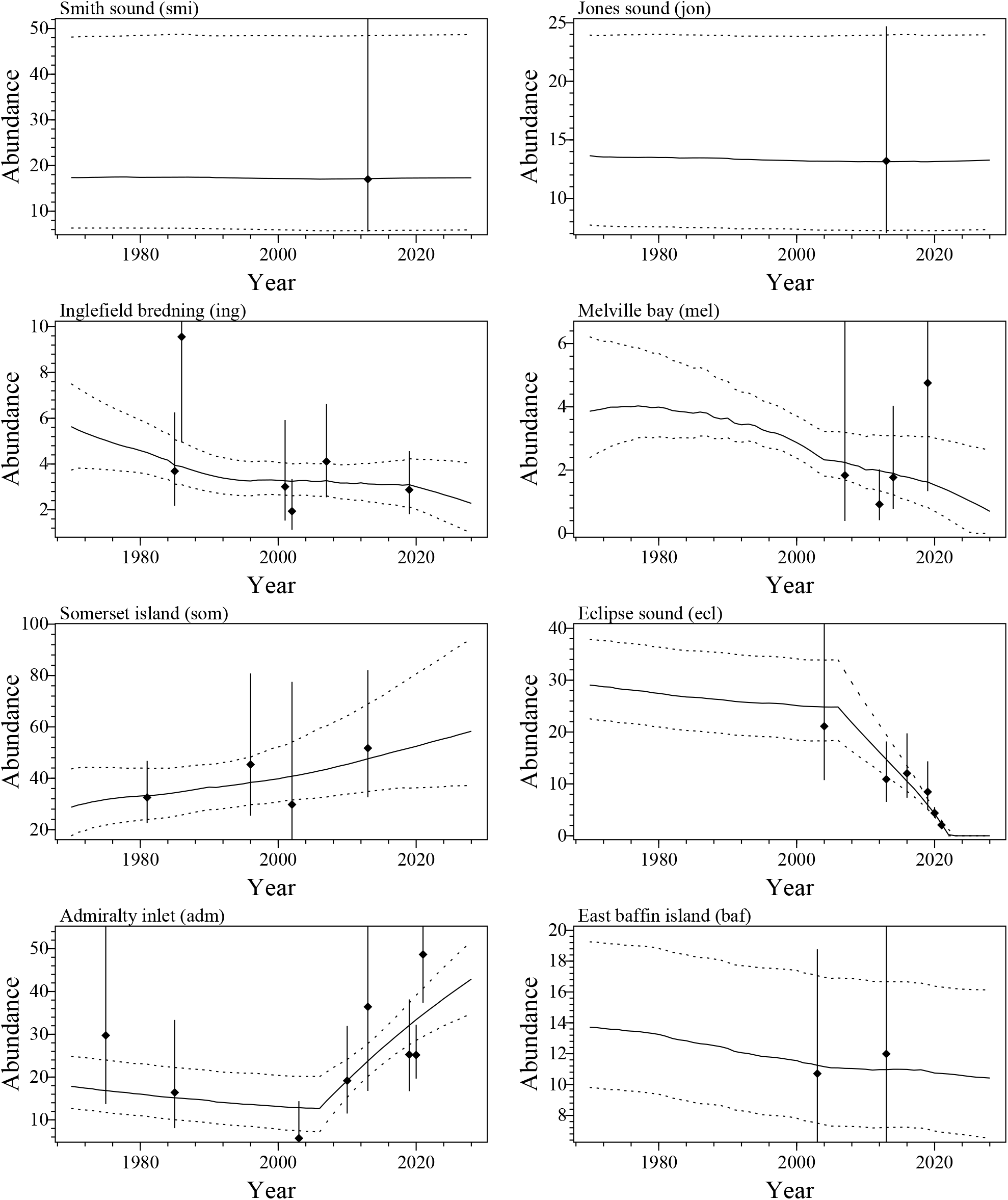
Estimated trajectories for narwhals in all areas. Points with bars are abundance estimates with 90% CI, solid curves the median of the estimated trajectories and dotted curves the 90% CI.

